# The transcription factor bZIP11 acts antagonistically with trehalose 6-phosphate to inhibit shoot branching

**DOI:** 10.1101/2023.05.23.542007

**Authors:** A. M. Hellens, P. Kreisz, J. L. Humphreys, R. Feil, J.E. Lunn, W. Dröge-Laser, C. A. Beveridge, F. Fichtner, C. Weiste, F. F. Barbier

## Abstract

The ontogenetic regulation of shoot branching allows plants to adjust their architecture in accordance with the environment. This process is due to the regulation of axillary bud outgrowth into branches, which can be induced by increasing sugar availability to the buds through decapitation of the shoot tip. Different sugar signalling components have been identified in the induction of shoot branching. However, the molecular components that maintain bud dormancy in response to sugar starvation remain largely unknown. Here, we show at the genetic level that basic leucine zipper 11 (bZIP11), a transcription factor that plays important roles in response to sugar starvation in plants, inhibits shoot branching in *Arabidopsis thaliana*. Physiology experiments demonstrated that bZIP11 protein levels are decreased by decapitation. Molecular and genetic evidence suggests that bZIP11 acts in a negative feedback loop with trehalose 6-phosphate (Tre6P), a sugar signal that promotes shoot branching. Our data also suggest that the central energy sensor SUCROSE NON-FERMENTING 1 RELATED KINASE1 (SnRK1), alleviates the inhibitory effect of Tre6P on bZIP11 protein accumulation and inhibits shoot branching. Altogether, these data provide a working model that involves bZIP11, Tre6P and SnRK1 in the regulation of shoot branching.

---

Plant shoot branching is an extremely plastic developmental trait that enables plants to adjust their architecture in order to acclimate to specific environmental conditions and compete with neighbouring plants for light harvest ^1^. This developmental process is also determining for the number of fruits and seeds set per plant, thereby contributing to plant fitness. In addition, due to its impact on yields of several crops, shoot branching is a key target for crop improvement ^2,3^. In flowering plants, shoot branching is mainly driven by the outgrowth of axillary buds at the axil of leaves ^4,5^. This developmental process is regulated by a hormonal network notably involving auxin, cytokinins and strigolactones ^4,5^. Sugars also play important roles in the control of shoot branching ^6–10^ and it is likely that sugars are the first signal to trigger bud outgrowth in response to removal of the shoot tip (decapitation) ^10^, a treatment that triggers axillary buds to grow out ^11,12^.

Independently of their metabolic functions, sugars play signalling roles in plant growth and development ^13,14^. Different signalling pathways enable sugar signals to be sensed and transduced. Several sugar signalling pathways have been shown to be involved in the control of shoot branching, including the pathways mediated by HEXOKINASE1 ^15^, which plays a role in glucose sensing ^13,14,16^, and trehalose 6-phosphate (Tre6P) ^17,18^, a sugar-metabolite that specifically signals sucrose availability ^19^. Different signalling pathways contribute to mediate the impact of sugar or carbon starvation in plants ^13,14,20,21^. A comparative transcriptomic study in buds highlighted that bud dormancy in annual and perennial plants is underpinned by a carbon starvation signalling network ^22^. This prompted us to investigate at the genetic and physiological levels the involvement of specific well-established components of sugar starvation signalling during the regulation of shoot branching.

The transcription factor bZIP11 is thought to be an important integrator of sugar starvation in plants ^21,23,24^. Indeed, bZIP11 belongs to a group of bZIP transcription factors whose translation is induced under low energy conditions ^23,25,26^, leading to transcriptional changes of bZIP-regulated genes that enable plants to adjust their metabolism, growth and development to unfavourable conditions ^21,24,26^. Some evidence also suggests that bZIP11 may act downstream of Tre6P ^27^, which triggers bud outgrowth in response to sugar availability ^17,18^. This prompted us to investigate whether bZIP11 could be involved in shoot branching. To test this, a genetic approach was first undertaken. To avoid issues due to redundancy with bZIP2 and bZIP44, which are phylogenetically very close to bZIP11 and have redundant functions (Kreisz *et al*., companion paper), we generated a triple *bzip2/11/44* knockout line using CRISPR/Cas9 (Kreisz *et al*., companion paper). Fifteen days after bolting, *bzip2/11/44* knockout plants show an increased shoot branching phenotype, with on average 2.5 more primary rosette branches than WT plants (Fig. 1a and b).

**Figure 1.**
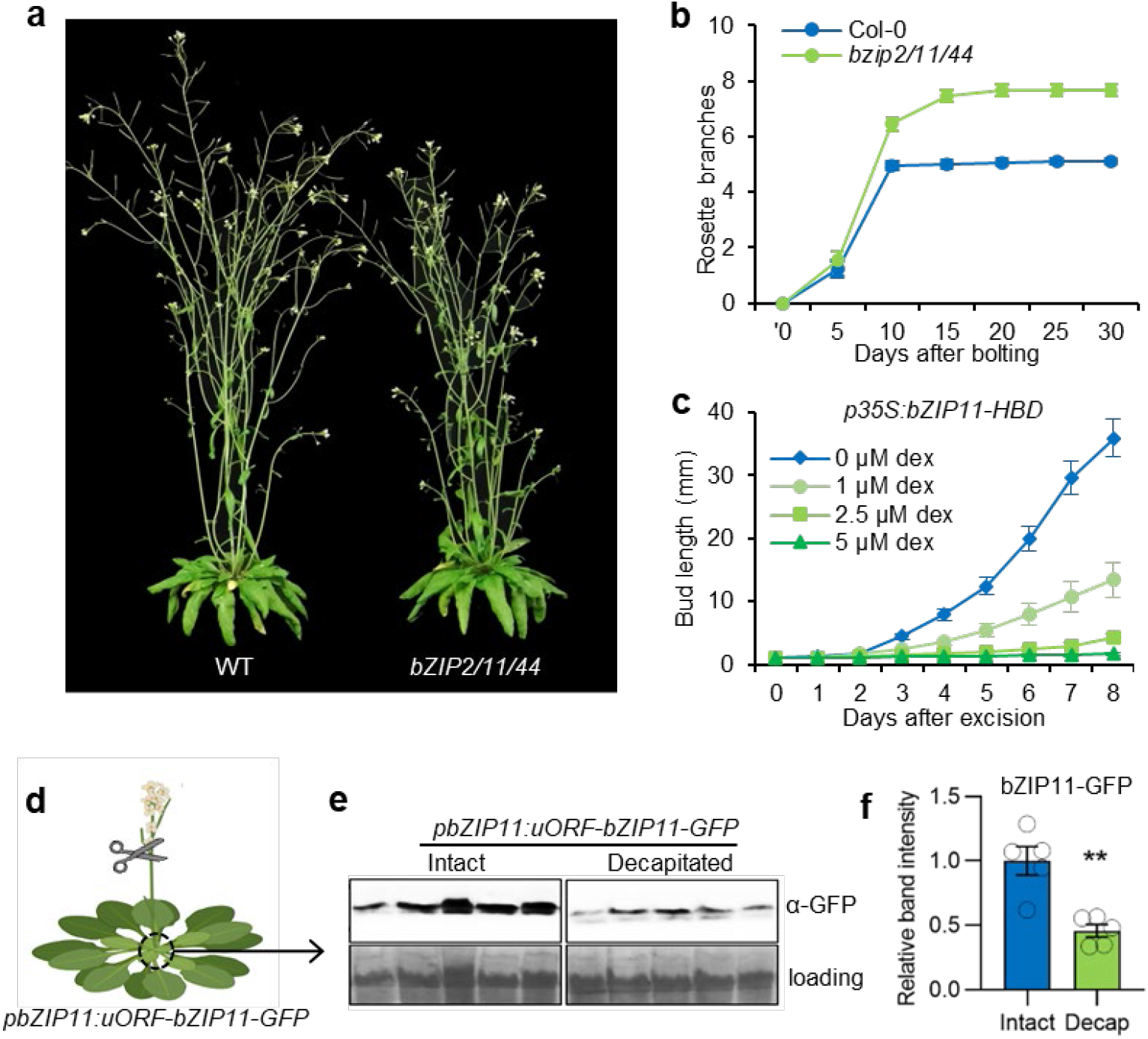
bZIP11 inhibits arabidopsis shoot branching. (**a**) Representative picture of 6-week-old plants and (**b**) number of primary rosette branches longer than 0.5 cm in WT (Col-0) and *bzip2/11/44* plants grown under 16 h photoperiod. Data are mean ± s.e.m (n = 20 plants). (**c**) Length of *p35S:bZIP11-HBD* single cauline buds grown on split plates with a range of dexamethasone (dex) in the growth media. Data are mean ± s.e.m (n = 12 buds). (**d**) Schematic representation of the decapitation assay performed in *Arabidopsis thaliana* ‘Columbia-0’ carrying a *pbZIP11:uORF-bZIP11-GFP* construct. (**e**) Western blot showing the accumulation of bZIP11-GFP in the core of arabidopsis rosettes in five individual intact and decapitated plants. Ponceau staining showing the Rubisco large subunit was used as a loading control. Average band intensity determined on the gel displayed in **e**, normalized by the loading control and relative to the intact plant conditions. Data are mean ± s.e.m (n = 5). Asterisks indicate the statistical significance (\*\*\**P*-value < 0.005).

The impact of bZIP11 on axillary bud outgrowth was then tested by inducing bZIP11 in cauline buds using an *in vitro* split-plate assay, commonly used to test the effect of different signals on bud outgrowth ^7,15,28^. To induce bZIP11, we used a previously published line in which bZIP11 translocation to the nucleus occurs in the presence of dexamethasone ^23^. Dexamethasone did not inhibit bud outgrowth in WT plants (Supp. Fig. S1), while dexamethasone-induction of bZIP11 inhibited bud outgrowth in a dose-dependent manner (Fig. 1c). Altogether, these results indicate that bZIP11, and likely its close homologues, play an inhibitory role during bud outgrowth and shoot branching in arabidopsis.

We then tested the involvement of bZIP11 in decapitation responses. To achieve this, the protein levels of bZIP11 were quantified in arabidopsis rosette cores, enriched in axillary buds, from decapitated and intact plants expressing a *pbZIP11:uORF-bZIP11-GFP* construct (Fig. 1c). The results show that 8 h after decapitation, bZIP11 protein level was strongly decreased when compared to intact plants (Fig. 1d-f), indicating that decapitation decreases bZIP11 protein levels. In addition, the *bzip2/11/44* mutant displayed a shoot branching pattern similar to the phenotype observed in decapitated WT plants (Supp. Fig. S2), supporting a role of bZIP11 in decapitation responses.

The role of bZIP11 in the control of bud outgrowth and shoot branching observed in Figure 1 is in contrast with the reported roles of Tre6P in these processes ^17,18^. This prompted us to investigate the potential connections between these two signalling pathways. As a previous report based on gene expression data of putative bZIP11 targets suggested that Tre6P may inhibit bZIP11 ^27^, we tested whether Tre6P could inhibit the accumulation of bZIP11 at the protein level. To test this, we transiently co-transfected arabidopsis leaf protoplasts with a *p35S:bZIP11-HA* and a *p35S:otsA* construct, which increases Tre6P levels by over-expressing the *Escherichia coli* TPS ^30,31^, and compared these with protoplasts transfected with a *p35S:GFP* construct used as a control (Fig. 2a). Western blot analysis indicates that the bZIP11 protein level was 25 times lower when co-transfected with *p35S:otsA* construct than with the *p35S:GFP* construct (Fig. 2a and b), showing that Tre6P accumulation strongly decreases bZIP11 protein levels in this system, thereby mimicking the effect of decapitation on bZIP11 protein levels.

**Figure 2.**
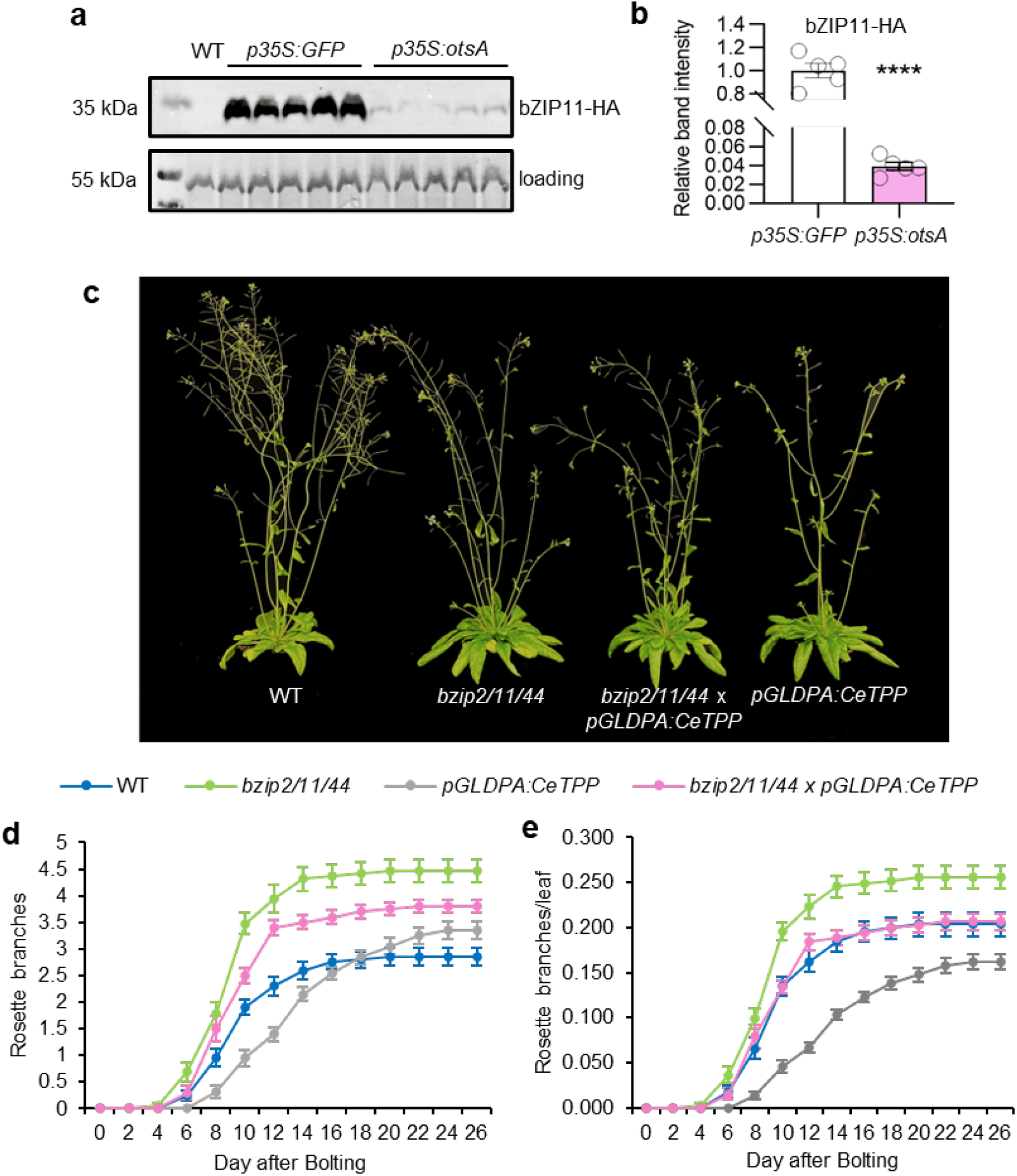
Tre6P inhibit bZIP11 accumulation in *Arabidopsis thaliana*. (**a**) Western blot showing ZIP11-HA accumulation in WT (Col-0) protoplasts co-transfected with *p35S:bZIP11-HA* plasmid and either *p35S:GFP* or *p35S:otsA* plasmids. Ponceau staining showing the Rubisco large subunit was used as a loading control. (**b**) Average band intensity determined on the gel displayed in **a**, normalized by the loading control. Data are mean ± s.e.m (n = 5). Asterisks indicate statistically significant differences determined by Student’s T-test (**** *P*-value < 0.0001). (**c**) Representative picture of 6-week old WT (Col-0), *bzip2/11/44, pGLDPA:CeTPP* and *bzip2/11/44* x *pGLDPA:CeTPP* plants grown under 16 h photoperiod. (**d**) Number of primary rosette branches longer than 0.5 cm and (**e**) number of primary branches divided by the number of rosette leaves. In **d** and **e**, data are mean ± s.e.m (n = 19-20 plants).

To then test whether this observation obtained in protoplasts might be relevant to shoot branching, we crossed the *bzip2/11/44* mutant with a *pGLDPA:CeTPP* line, a construct that decreases Tre6P levels by overexpressing a *TREHALOSE 6-PHOSPHATE PHOSPHATASE* (*TPP*) from *Caenorhabditis elegans* in the vasculature and which was previously reported to inhibit shoot branching ^17^. In our conditions, the *pGLDPA:CeTPP* line only had a transient negative effect on the number of rosette branches produced (Fig. 2c and d), while the *bzip2/11/44* mutant produced more branches than the WT, as observed in Figure 1a and b. The shoot branching phenotype of the *bzip2/11/44* x *pGLDPA:CeTPP* cross was intermediate between the *bzip2/11/44* and *pGLDPA:CeTPP* line (Fig. 1c and d). It was previously reported that the number of nodes produced by the rosette may affect the number of rosette branches produced ^29^. As these lines have delayed flowering and produce more nodes than the WT (Supp. Fig. S3), we divided the number of rosette branches by the number of rosette leaves to normalise the branching data. This quantification method revealed, in comparison to WT, a stronger decrease in the shoot branching phenotype of the *pGLDPA:CeTPP* line (Fig. 2e), while the *bzip2/11/44* mutant retained an increased shoot branching phenotype. Under this analysis, the *bzip2/11/44* x *pGLDPA:CeTPP* cross had an intermediate phenotype, in this case, similar to the WT phenotype (Fig. 2e). The results of this phenotypic analysis show that knocking out bZIP11 and its close homologues compensates part of the decreased branching phenotype observed in the *pGLDPA:CeTPP* line.

The data presented in Figure 2 also indicate that decreasing Tre6P levels by introducing the *pGLDPA:CeTPP* construct in the *bzip2/11/44* mutant inhibits part of the branching phenotype observed in this mutant. A previous study suggested that bZIP11 might induce Tre6P dephosphorylation through induction of the expression of *TREHALOSE 6-PHOSPHATE PHOSPHATASE* (*TPP*) genes ^24^. We therefore tested whether part of the branching phenotype of the *bzip2/11/44* mutant could be due the lack of bZIP11 promoting Tre6P dephosphorylation. Using the same protoplast system as in Fig. 2a, we identified by qRT-PCR five *TPP* genes with significantly increased expression in response to bZIP11 induction (*TPPB, TTD, TPPE*, TPPF and *TPPH;* Fig 3a). Among these, *TPPF* was previously reported to be induced by bZIP11 ^24^. DNA affinity purification with sequencing (DAP-seq) data ^30^ were then used to assess whether the bZIP11 transcription factor could directly target these five *TPP* genes *in vitro*. Visualization of the DAP-seq data indicated that bZIP11 can bind to the promoter of all five *TPP* genes identified as induced by bZIP11 in this study (Fig. 3b).

**Figure 3.**
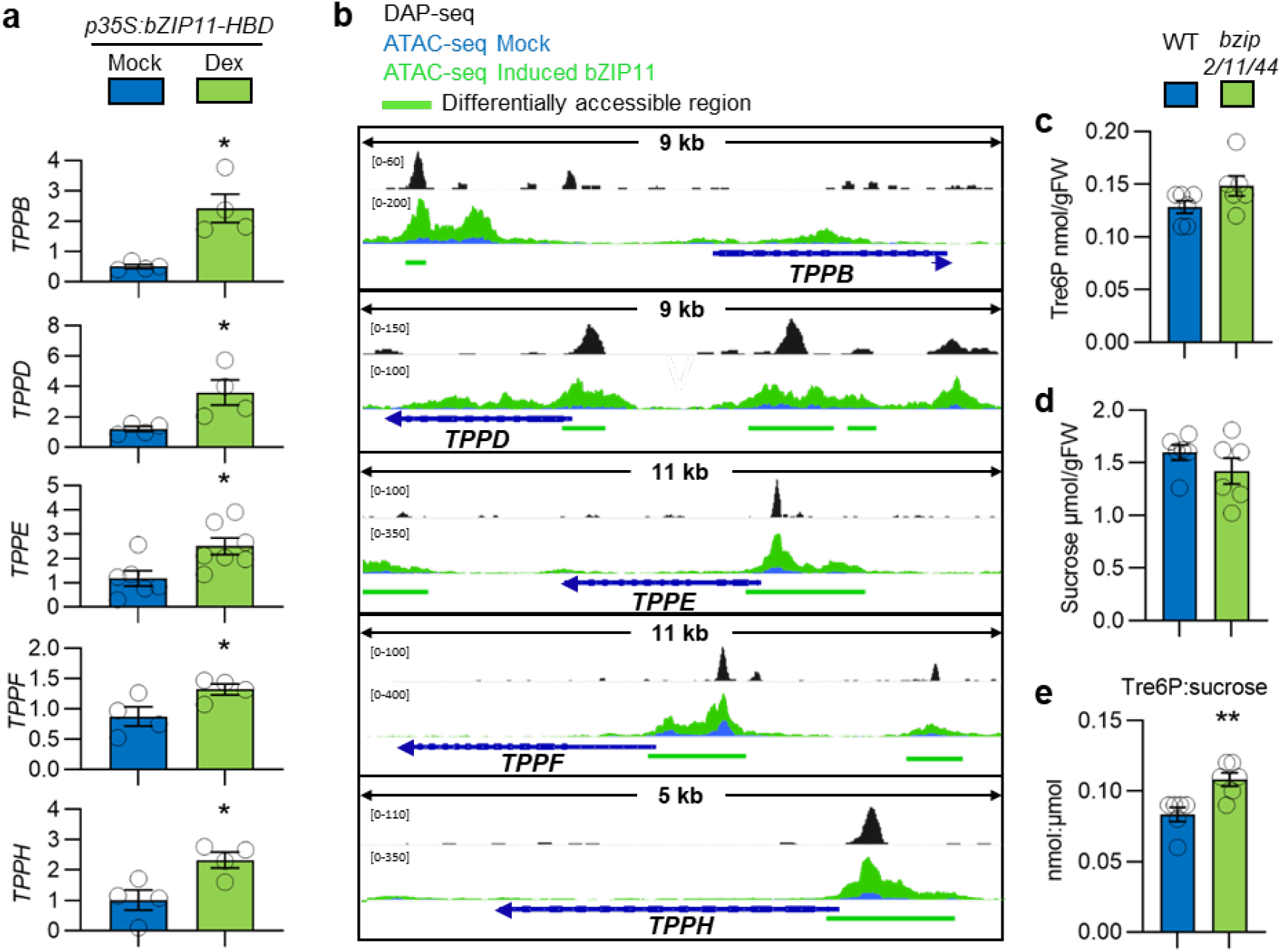
bZIP11 promotes Tre6P dephosphorylation in *Arabidopsis thaliana*. (**a**) Expression of *TREHALOSE-6-PHOSPHATE PHOSPHATASE* (*TPP*) genes significantly regulated in response to bZIP11 induced with 10 µM dexamethasone applied to *p35S:bZIP11-HBD* protoplast for 45 min. (**b**) Genome browser view showing DNA Affinity Purification (DAP)-seq performed with bZIP11 and Assay for Transposase Accessible Chromatin (ATAC)-seq signals around *TPP* genes significantly regulated upon bZIP11 induction as described in **a**. (**c**) Tre6P levels, (**d**) sucrose levels, and (**e**) Tre6P to sucrose ratio measured in WT (Col-0) and *bzip2/11/44* mutant plants. Whole 4-week-old rosettes were harvested at ZT6. Data are mean ± s.e.m (n = 6). (**d**) In **c** and **e**, asterisks indicate statistically significant differences determined by Student’s T-Test (** *P*-value <0.001, **** *P*-value < 0.001).

As bZIP11 was reported to promote gene expression by modifying chromatin accessibility ^31^, we tested whether bZIP11 could regulate chromatin accessibility around *TPP* genes that are induced by this transcription factor. To assess whether bZIP11 could regulate chromatin accessibility around these five *TPP* genes we performed an Assay for Transposase Accessible Chromatin with next-generation sequencing (ATAC-seq) after induction of bZIP11 in arabidopsis protoplasts. After 45 min of bZIP11 induction, we observed differentially accessible regions (DARs) of chromatin upstream of these five *TPP* genes (Fig. 3b). The positions of the DARs overlap with the positions of the DAP-seq peaks (Fig. 3b), indicating that the changes in chromatin accessibility may be due to direct binding of bZIP11 to the five *TPP* loci.

Evidence that bZIP11 may negatively regulate Tre6P levels in arabidopsis is supported by a previous study showing that bZIP11 induction decreases Tre6P levels in this species ^24^. We therefore tested whether the *bzip2/11/44* mutant accumulates more Tre6P than the WT. Whole-rosette measurements of Tre6P and sucrose levels showed no significant differences between the *bzip2/11/44* and the WT (Fig. 3c and d), but the Tre6P:sucrose ratio was significantly higher (1.3-fold) in the *bzip2/11/44* plants than in the WT (Fig. 3e), supporting the hypothesis that bZIP11 negatively regulates Tre6P levels in arabidopsis, likely by enhancing Tre6P dephosphorylation through TPPs (Fig. 3a and b) ^24^.

Tre6P has been suggested to act, at least partly, through inhibiting the activity of the SUCROSE NON-FERMENTING 1 RELATED KINASE 1 (SnRK1) complex via either direct or indirect binding to the SnRK1α1 catalytic subunit ^32,33^. SnRK1 is a master regulator of energy homeostasis that is activated under starvation conditions ^14,20,34^ and induces the activity of the S_1_/C bZIPs. SnRK1 phosphorylates bZIP63 (a group C bZIP) at three specific serine residues ^35^, which triggers preferential heterodimerization with the group S_1_ bZIPs rather than homodimerization with group C bZIPs, a mechanism that modifies their transactivation properties and reprograms their targets ^21,35,36^. Given the connections among bZIP11, Tre6P, and SnRK1, we investigated whether increasing SnRK1 activity could alleviate the inhibitory effect of Tre6P on bZIP11 accumulation (Fig. 2d). To achieve this, we co-transfected arabidopsis leaf protoplasts with a *p35S:bZIP11-HA* and a *p35S:otsA* construct, as in Fig. 2a, but this time, we also co-transfected an increasing amount of *p35S:SnRK1α1* construct (Fig 4a and b). Western blot analysis of bZIP11-HA protein levels showed that the inhibitory effect of Tre6P on bZIP11 accumulation is gradually alleviated by increasing the amount of *p35S:SnRK1α1* construct co-transfected (Fig. 4a and b). In addition, in absence of *p35S:otsA*, the *p35S:SnRK1α1* led to increased accumulation of bZIP11 (Fig. 4a and b). These results indicate that *SnRK1α1* activity promotes bZIP11 protein levels and alleviates the inhibitory effect of Tre6P. This also suggests that the positive effect of SnRK1 on the S_1_/C bZIPs is not limited to promoting their heterodimerization and that SnRK1 also contributes to increasing the protein levels of bZIP11.

**Figure 4.**
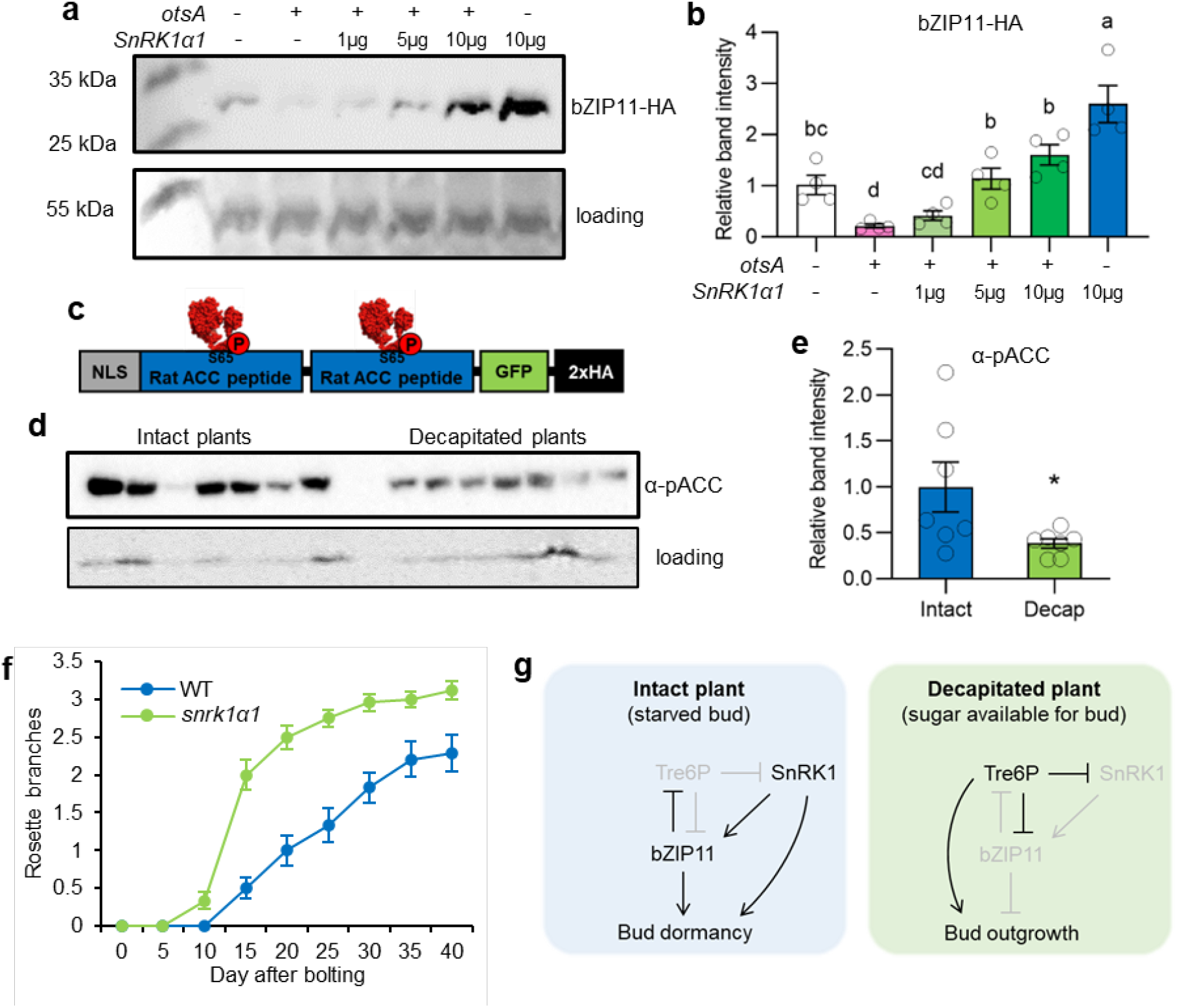
SnRK1 promotes bZIP11 accumulation in leaf protoplasts and inhibits shoot branching in *Arabidopsis thaliana*. (**a**) Western blot showing bZIP11-HA accumulation in WT (Col-0) protoplasts co-transfected with *p35S:bZIP11-HA* and different concentration of *p35S:otsA* and *p35S:SnRK1α1* plasmids as indicated above the blot. Ponceau staining was used as a loading control. (**b**) Average band intensity determined on the gel displayed in **a**, normalized by the loading control. Data are mean ± s.e.m Statistical differences are indicated by one-way AVOVA no multiple correction (Fishers LSD test). (**c**) Representation of the *p35S:NLS-ratACC-GFP-HA* construct used to assess SnRK1 activity. Using a pYX242 expression vector, an AMP-activated protein kinase (AMPK, shown in red) phosphorylated rat ACC peptide (blue) is expressed. (**d**) SnRK1 activity in five-week-old intact plants or plants decapitated for 8 h and harbouring the construct shown in **c**. Unspecific band was used as a loading control. (**e**) Average band intensity determined on the gel displayed in **d**, normalized by the loading control and relative to the intact plant conditions. Data are mean ± s.e.m (n = 7) and the asterisks indicate the statistical significance (\**P*-value < 0.05). (**f**) Number of primary rosette branches longer than 0.5 cm in WT (Col-0) and *snrk1α1* plants grown under 16 h photoperiod. Data are mean ± s.e.m (n = 20 plants). **(g)** Working model showing the role of bZIP11 in the regulation of bud outgrowth as described in the text (the lines do not necessarily represent direct connections).

To then explore whether SnRK1 was likely involved in the regulation of shoot branching, we assessed the response of SnRK1 activity to decapitation using a *p35S:NLS-ratACC-GFP-HA* reporter line. This line harbours the rat ACETYL-COA CARBOXYLASE (ACC), which is a target of the mammalian SnRK1 ortholog, AMPK ^37,38^ (Fig. 4c). The phosphorylation status of ACC determined through Western blot analysis is used as a proxy for SnRK1 activity. The results show that SnRK1 activity was 2.5 times lower in rosette cores six hours following decapitation (Fig. 4d and e), suggesting that reduced SnRK1 activity is associated with decapitation-induced shoot branching. To verify this, we determined the shoot branching phenotype of the *snrk1α1* KO mutant ^36^. Phenotypic analysis of *snrk1α1* revealed that this mutant produced significantly more primary rosette branches than the WT (Fig. 4f), supporting the hypothesis that SnRK1 negatively regulates shoot branching.

In conclusion, the evidence presented in this study allows us to propose a working model concerning the involvement of molecular components of sugar starvation signalling that contribute to the control of shoot branching (Fig. 4g). In this model, we propose that in intact plants, where axillary buds are carbon starved, the transcription factor bZIP11 inhibits axillary bud outgrowth and shoot branching, presumably via enhancing degradation of Tre6P by TPPs (Fig. 2 a-e), a sugar signal that reflects sucrose availability ^19^. In decapitated plants, where more sugars are available for axillary buds, Tre6P increases in axillary buds ^17,18^ and decreases bZIP11 protein levels (Fig. 2a), thereby triggering bud outgrowth. Our data also suggest the involvement of the energy sensor kinase complex SnRK1 ^13,34,39^ in this process. In intact plants, the activity of SnRK1 stays high, keeping bZIP11 protein levels high and maintaining bud dormancy. In decapitated plants, SnRK1 activity is inhibited, allowing Tre6P to inhibit bZIP11 and promote bud outgrowth. Based on this working model, we propose that the balance between bZIP11 and Tre6P acts as a homeostatic mechanism to modulate shoot branching in response to changes in sugar availability and allocation triggered by decapitation or environmental factors. These discoveries should prompt future research to test whether the working model proposed here concerns other developmental processes. In addition, given the strong impact of shoot branching on crop yields and the potential of bZIP11, Tre6P and SnRK1 for improving crops ^13,40^, this study should increase the toolkit of research programs aimed at improving crops.

## MATERIAL AND METHODS

### Plant material and growth conditions

All *Arabidopsis thaliana* plants are in the Columbia-0 background. *pbZIP11:bZIP11-GFP, p35S:bZIP11-HBD-M* ^23^, *snrk1α1* ^36^, *p35S:NLS-ratACC-GFP-HA* ^37^, and *pGLDPA:CeTPP*.*4* ^17^ plants have been described earlier. *bzip2/11/44* mutants were produced by CRISPR/Cas9 system ^41^. Target specific sequences were selected using ChopChop ^42^. Guides were positioned close to the ATG. Homozygous lines were selected with a single bp insertion, resulting in frameshift and premature stop.

Arabidopsis seeds were stratified for 3 d at 4°C then transferred to growth chambers with 16 h:8 h, light : dark, 22:20°C, day : night with either low (70 ± 10 µmol m^−2^ s^−1^) or high (150 ± 20 µmol m^−2^ s^−1^) light intensity as indicated by each individual experiment. Plants were grown in UQ23 potting mix supplemented with dolomite and osmocote or Soil SP Pikier from Gebr. Patzer GmbH & Co. Decapitation was performed on 5-week-old plants, approximately 10 d after bolting and involved removal of the shoot tip 10 cm above the rosette and removal of any emerged cauline branches. Bud-enriched material was harvested by removal of all the leaves, upper stem, and hypocotyl from the core of the rosette.

### RNA extraction and gene expression

Total RNA for real-time quantitative PCR (RT-qPCR) was extracted either using a NucleoSpin RNA, Mini kit (Macherey-Nagel), or as described in ^43^. Briefly, ground samples were lysed for 15 min in a CTAB/PVP buffer supplemented with DTT to prevent RNA degradation. Nucleic acids were precipitated in isopropanol and pelleted by centrifugation for 45 min at 20,000g. Ethanol-washed pellets were resuspended in water and a DNase treatment was applied for 25 min at 37°C. Total RNA was then precipitated in isopropanol and pelleted by centrifugation for 45 min at 20,000g. RNA was then eluted in water and the quality of the RNA was assessed by electrophoresis.

RNA was then converted into cDNA by reverse transcription flowing the manufacturer’s instructions (iScript Supermix, Bio-Rad Laboratories, California, USA). The diluted cDNA was then used as a template for quantitative Real-Time PCR following the manufacturer’s instructions (SensiFAST™ SYBR® No-ROX Kit; Bioline). Samples were amplified following the manufacturer’s instructions and fluorescence was monitored with a CFX384 Touch™ Real-Time PCR Detection System (Bio-Rad Laboratories, California, USA) using the following protocol: 3 min 95°C, 40 cycles at 10 s 95°C, 45 s at 59°C, and 1 min 95°C, 1 min 55°C). Gene expression was calculated using the ΔΔCt method and corrected by primer efficiency. Gene expression was normalized to the average of two technical replicates and geomean expression of *ACTIN* (Combination of *ACT2, ACT7*, and *ACT8*: *At3Gg18780, At5g09810, At1g49240*), *TUBULIN3* (*At5g62700*) and *18S* (*18S rRNA*). All primer sequences used in this paper are detailed in Extended Data Table 1.

### Vectors and cloning

*p35S:otsA* ^44^, *p35S:bZIP11-HA, p35S:GFP, p35S:SnRK1α1-HA* ^46^ constructs have been previously described.

### Branching measurements

Primary rosette branches longer than 5 mm in length were counted every 2 or 5 days (depending on individual experiment) after bolting.

### Split plate assay

Single stem segments containing a single unexpanded node were excised from cauline stems. WT and *p35S:bZIP11-HBD* plants were used for this experiment. Stem segments were placed on half-strength MS media supplemented with 30 mM sucrose and either 0 μM, 1 μM, 2.5 μM, or 5 μM of dex. Plates contained 12 individual stems of each genotype per treatment. Plates were placed back in growth chambers vertically and monitored daily for eight days. Bud length was determined by analysing photographs of buds using ImageJ.

### Transient expression assay in protoplasts

Protoplast isolation and transfection was carried out as described in ^47,48^. Plasmid DNA was transformed into *Escherichia coli* and overnight cultures were purified using pDNA Midiprep (Thermo Fisher Scientific). Plasmid DNA was prepared in a total of 20 μL per reaction at 1 to 40 μg concentrations of the various plasmid combinations as detailed for each experiment. *p35S:GFP* was always used as a control and where different concentrations of *p35S:otsA* and *p35S:KIN10* were used in Fig. 4a DNA concentration was made up to a total of 40 μg with *p35S:GFP*. Arabidopsis leaf protoplasts were extracted from 4-week-old plants by placing 0.5-1 mm cuts perpendicular to the midrib on abaxial side of the leaf and placing ∼30 leaves abaxial side down in 10 mL of enzyme solution (1 % Cellulase ‘Onozuka’ R10, 0.25 % maceroenzyme ‘Onozuka’ R10, 0.4 M mannitol, 20 mM KCl, 20 mM MES, 10 mM CaCl_2_, 0.1 % BSA, adjust to pH 5.7). Vacuum infiltration was applied to the leaves for 1 h then left at room temperature for a further 3 h to continue digestion. Digested cells were filtered through 75 μM mesh, then washed twice in ice-cold W5 (154 mM NaCl, 125 mM CaCl_2_, 5 mM KCl, 2 mM MES). Protoplasts were then left on ice for 1 h to settle at the bottom. Protoplasts were resuspended in MMg (0.4 M mannitol, 15 mM MgCl_2_, 4 mM MES, adjusted to pH 5.7) and kept at room temperature. 200 μL of protoplasts were added to each plasmid DNA reaction and tubes are inverted gently several times to mix cells with DNA. 220 μL of 40 % PEG was added to each tube and inverted several times to mix then incubated at room temperature for 20 mins. Protoplasts were pelleted then re-suspended in 300 μL of Wi solution (0.5 M mannitol, 2 mM KCl, 4 mM MES). Transfection reactions took place overnight for 16 h in growth chamber conditions. Supernatant was removed and cells were frozen in liquid nitrogen.

### SDS-PAGE Western Blot

For protoplast transformations, cells were snap frozen in liquid nitrogen then 150 μL of protein extraction buffer (4M urea, 16.6 % glycerol (v/v), 5 % ß-mercaptoethanol, 5 % SDS, and bromophenol blue) was added to a pool of three separate transformations. For plant tissue, protein extraction buffer was added to ground, frozen samples at a 2:1 volume:weight ratio. Lysate was then vortexed and boiled at 70°C for 10 min. 15 μL of lysate was loaded into an individual well of a 12 % polyacrylamide gel. Proteins were separated by Sodium dodecyl sulphate-polyacrylamide gel electrophoresis at a voltage of 100 V. Proteins were then transferred to PVDF membrane (Immobilon, Millipore, Billerica, MA, USA) by semi-dry blotting. Membranes were blocked for 1 h in 2 % skim milk. GFP-tagged proteins were detected by HRP-coupled anti-GFP at a dilution of 1:2000, with a secondary goat-anti-rat IgG-HPR at a dilution of 1:3000 (Chromo Tek [3H9] and [SA00004-8]). HA-tagged proteins were detected by HRP-coupled anti-HA at a dilution of 1:2000, with a secondary goat-anti-rabbit IgG-HPR at a dilution of 1:3000 (ChromoTek [7C9] and [SA00001-2]. ACC was detected by phospho-Acetyl-CoA Carboxylase (Ser79) Antibody (Cell Signaling Technology, Danvers, MA, USA). All blots were incubated for each antibody at 4°C overnight and all antibodies were diluted in 2 % skim milk. imaged using enhanced chemiluminescence (Clarity and ChemiDoc, Bio-Rad Laboratories, California, USA). Semi-quantitative band density was determined using ImageJ and normalised by Ponceau staining for whole protein loading control.

### Protoplast extraction for ATAC-seq

Four-week-old WT and *p35S:bZIP11-HBD* plants were used to extract mesophyll protoplasts via the epidermal leaf peel method (F.-H. Wu et al., 2009). Six leaves were placed in 10 mL of enzyme solution (solutions used here are the same as in the previous section) and digested for 1 h with constant gentle agitation. Cells were filtered with 50 μM mesh (CellTrics, Sysmex, Norderstedt, Germany) then washed twice in W5. Protoplasts were then re-suspended in MMg at a concentration of 200,000 cells per mL. Reactions took place in 2 mL in six-well plates with constant agitation. WT cells were treated with 10 μM dex dissolved in 100 % acetone, mock plants were treated with acetone, and *p35S:bZIP11-HBD* plants treated with 10 μM dex. 45 min after treatment, cells were spun down and split, taking 500 μL for ATAC-seq and 1.5 mL for RNA extraction.

### ATAC-seq library preparation

ATAC-seq library preparation was performed as modified from ^45,46^. Following protoplast extraction, nuclei were isolated from approximately 50,000 cells per reaction by sucrose sedimentation, modified from ^47^. Freshly extracted cells were centrifuged at 500 x g at 4°C. The following steps were all carried out on ice. Supernatant was discarded and pellet was resuspended in 1 mL of ice-cold nuclei purification buffer (20 mM MOPS, 40 mM NaCl, 90 mM KCl, 2 mM EDTA, 0.5 mM EGTA, 0.5 mM spermidine, 0.2 mM spermine 1 x protease inhibitors, adjust to pH 7). Cells were then filtered through 30 μM mesh (CellTrics, Sysmex, Norderstedt, Germany). Nuclei were then spun down at 1200 x g for 10 min at 4°C and pellet was resuspended in 1 mL of ice-cold nuclei extraction buffer 2 (0.25 m sucrose, 10 mM Tris-HCl pH8, 10 mM MgCl, 1 % Triton X-100, 1 x protease inhibitors). This step was repeated but this time pellet was resuspended in 300 μL of NPB and this resuspension of nuclei was carefully layered over 300 μL of ice-cold nuclei extraction buffer 3 (1.7 M sucrose, 10 mM Tris-HCl pH 8, 2 mM MgCl, 0.15 % Triton X-100 1 x protease inhibitor). The two layers were then spun down at 300 x g for 20 min at 4°C following which the supernatant was removed. Nuclei were resuspended in 50 μL of tagmentation reaction mix as per manufacturer instructions (TDE1, Illumina) and incubated at 37°C for 30 mins with gentle agitation every 5 min. Reactions were purified following manufacturer’s instructions using a QIAGEN MinElute PCR purification kit (catalogue number 28004) and eluted in 11 μL of elution buffer. DNA was amplified by PCR using ATAC barcoded primers and NEB Next High-Fidelity PCR Master Mix (5 min 72°C, 30 sec 98°C, then 5 x (10 sec 98°C, 30 sec 63°C, 1 min 72°C) held at 4°C). 5 μL of the PCR reaction was then further amplified by qPCR (30 sec 98°C, then 20 x (10 sec 98°C, 30 sec 63°C, 1 min 72°C)) to determine the required number of additional cycles. Additional cycle number for each reaction was determined by the cycle number for which a reaction has reached one third of its maximum, using the linear fluorescence vs cycle number graph from the qPCR. All libraries were purified with AMPure XP beads at a ratio of 1.5 : 1 beads : PCR reaction. Final elution in 20 μL of 10 mM Tris pH 8. Libraries were sequenced using Illumina HiSeq paired end 150 bp by NovoGene, Singapore.

### ATAC-seq processing and identification of DARs

Processing was carried out using Galaxy Australia (The Galaxy Community, 2022) and R with RStudio (Version 4.2.2) with the following steps. In Galaxy, raw reads were trimmed using Trimmomatic ^48^ with a 10 bp HEADCROP, a SLIDINGWINDOW with an average quality of 30 over every 6 bp, and an ILLUMINACLIP NexteraPE. Reads shorter than Reads were mapped against Arabidopsis thaliana TAIR10 reference genome using Bowtie2 ^49^, with paired end, dovetailing, and a maximum fragment length of 1000. Reads smaller than 30 bp, duplicate reads, reads with a quality score of <30 phred, and those which were mapped to the chloroplast or mitochondrial genome were discarded. Peaks were called with MACS2 ^50^ using the inputs: single-end BED, effective genome size 1.2e8, an extension size of 200 and a shift size of 100. BED and BAM and index files were then imported into RStudio and DARs were determined using the package DiffBind ^51^. Peaks were read with peakCaller=“narrow”, minOverlap=3 and dba.contrast function was specified to compare mock treated and bZIP11-induced samples. The package rtracklayer was used to convert the DiffBind peaks report into BED format. The peaks report was then imported back into Galaxy where differential peaks were annotated to the Arabidopsis TAIR10 reference genome using ChIPseeker ^52^. The resulting BED file of annotated DARs was imported into the Interactive Genome Viewer (Broad Institute, University of California) ^53^ along with BED files of samples from MACS2 output for visualisation.

### Sucrose and Tre6P measurements

Four-week-old whole rosettes were frozen in liquid nitrogen and ground into a fine powder. Water-soluble components were extracted as described in ^54^. Sucrose was measured spectrophotometrically by sequential enzymatic reactions ^55^. Tre6P levels were determined as described in ^54^ with modifications as described in ^56^.

## ACKNOWLEDGMENT

Christine Beveridge is the recipient of an Australian Research Council Centre of Excellence CE200100015 and an Australian Laureate Fellowship FL180100139. Alicia Hellens is supported by a Commonwealth Scientific and Industrial Research Organization Postgraduate Scholarship. Philipp Kreisz was supported by a research grant from the German Research Foundation to CW (DFG project number: 408153945). Regina Feil and John Lunn were financially supported by the Max Planck Society. Figures 1c was made using BioRender.com.

## SUPPORTING INFORMATION

**Table S1.**
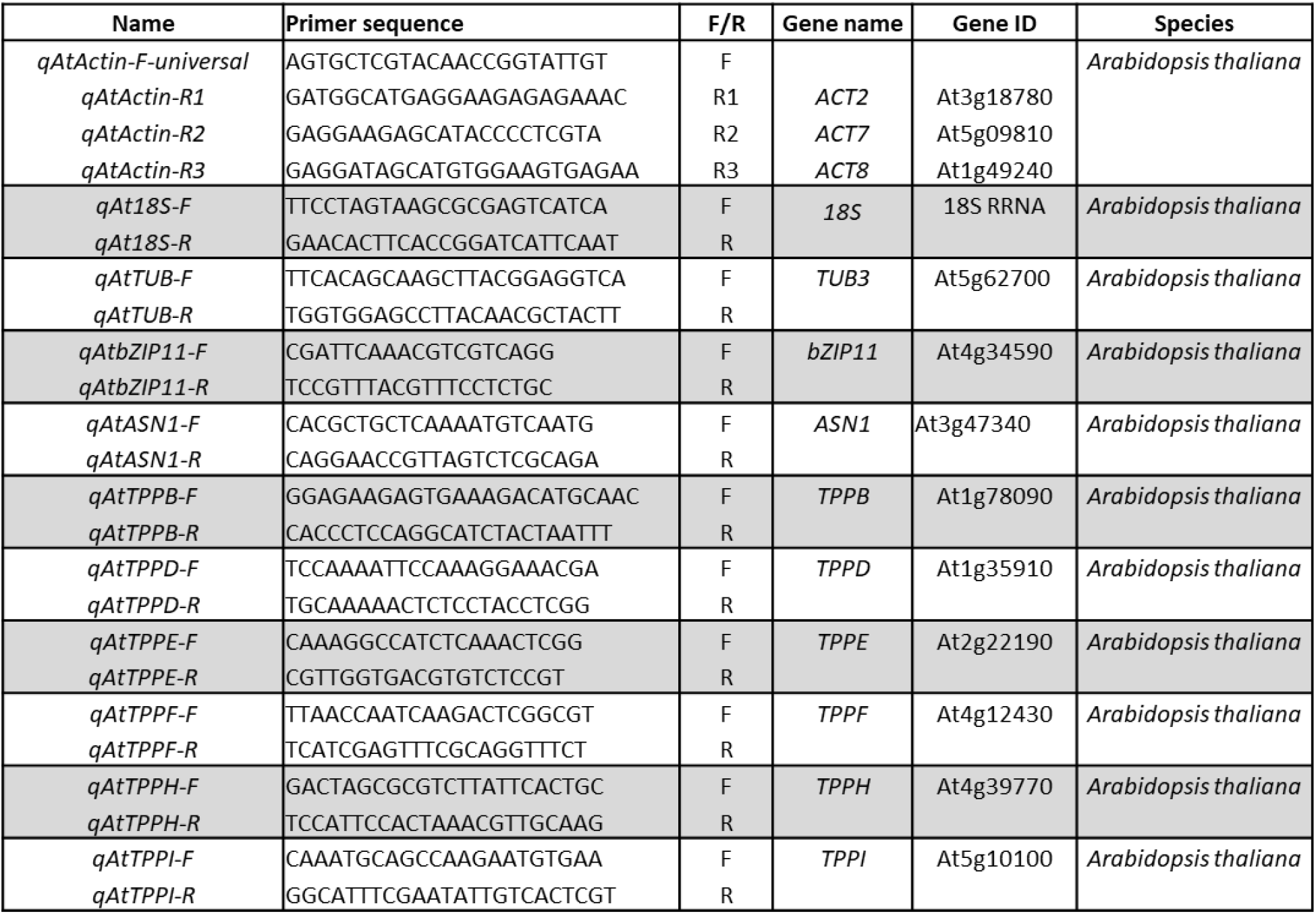
List of primers used in this study.

**Supplementary Figure S1.**
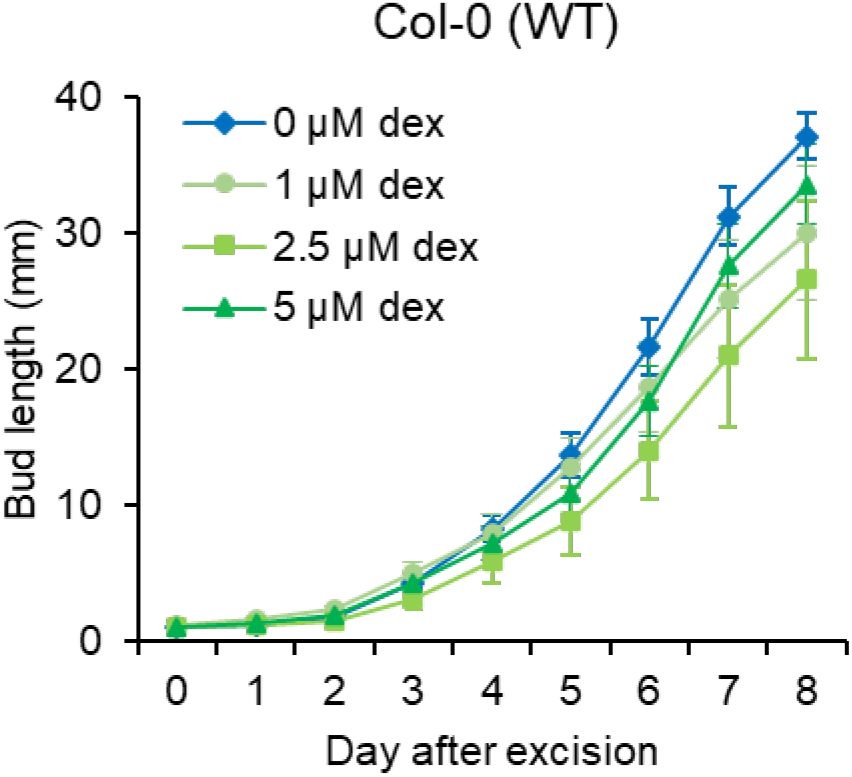
Impact of dexamethasone (dex) on cauline bud elongation in WT plants. Length of *p35S:bZIP11-HBD* single cauline buds grown on split plates with a range of dexamethasone (dex) in the growth media. Data are mean ± s.e.m (n = 12 buds).

**Supplementary Figure S2.**
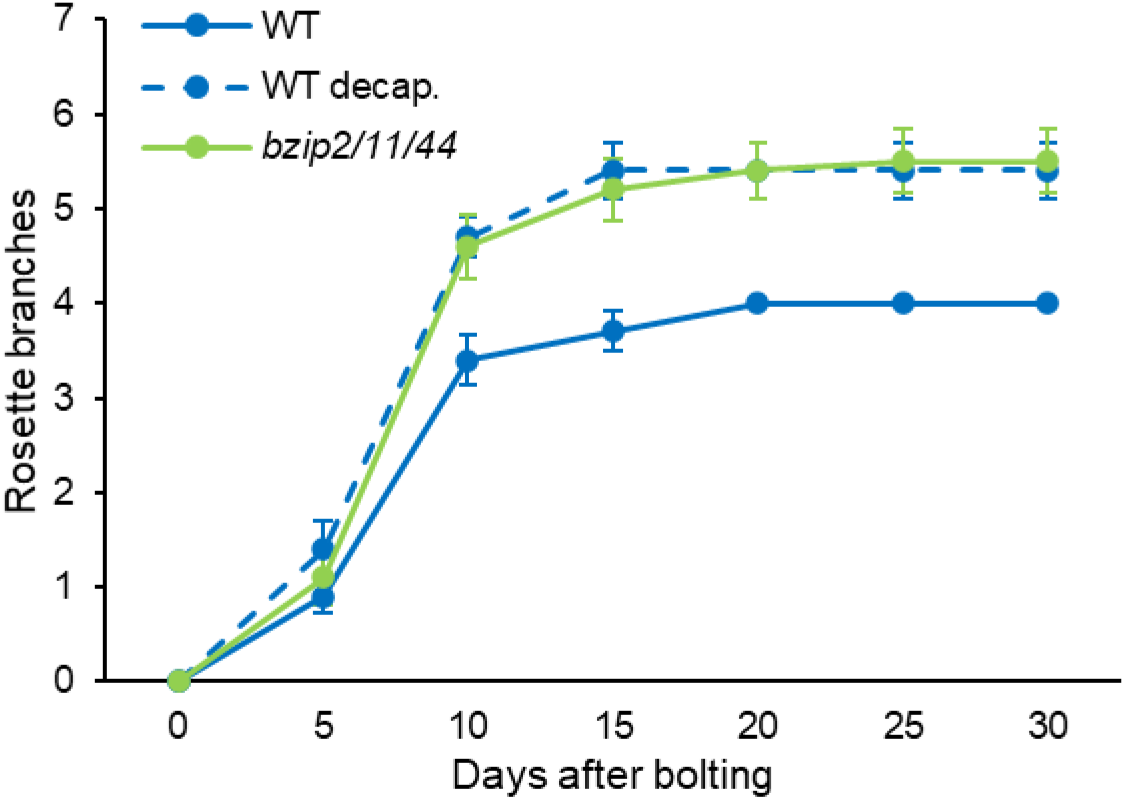
Shoot branching phenotype in *bzip2/11/44*, and decapitated WT. Number of primary rosette branches longer than 0.5 cm in WT (Col-0) intact and decapitated (decap.) and *bzip2/11/44* plants grown under 16 h photoperiod. Data are mean ± s.e.m (n = 20 plants).

**Supplementary Figure S3.**
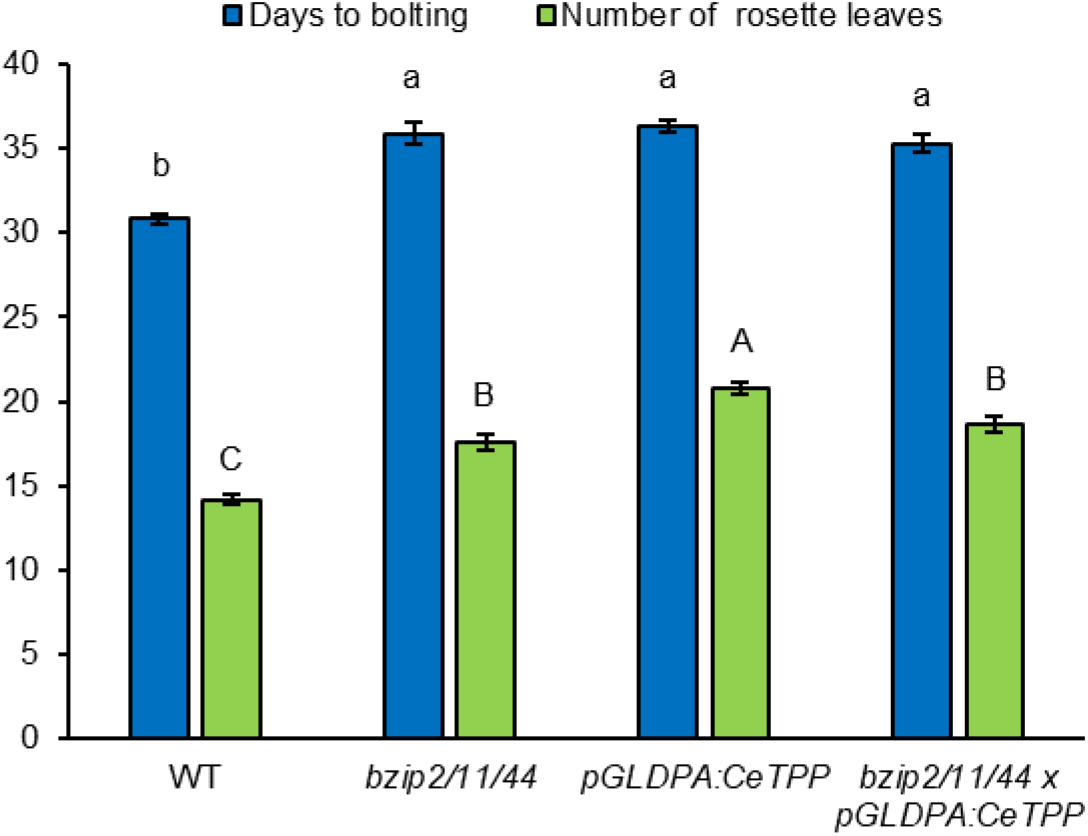
(a) Time to bolting and number of rosette leaves produced by the lines presented in figure 2. Data are mean ± s.e.m (n = 19-20 plants). Letters on the graph indicate statistical difference determined by two-way ANOVA with Šídák’s multiple comparisons test.

